# Herd Immunity to Ebolaviruses is not a Realistic Target for Current Vaccination Strategies

**DOI:** 10.1101/289249

**Authors:** Stuart G. Masterson, Leslie Lobel, Miles W. Carroll, Mark N. Wass, Martin Michaelis

**Affiliations:** Industrial Biotechnology Centre and School of Biosciences, University of Kent, Canterbury, UK; Department of Microbiology, Immunology and Genetics, Faculty of Health Sciences, Ben-Gurion University of the Negev, Beer-Sheva, Israel; Department of Emerging and Re-emerging Diseases and Special Pathogens Uganda Virus Research Institute (UVRI), Entebbe, Uganda; Public Health England, Porton Down, Salisbury, United Kingdom (M.W. Carroll)

**Author notes:** Correspondence: Mark N. Wass,; Martin Michaelis.

**Keywords:** Ebola virus, Ebolavirus, Vaccines, Herd immunity, Basic Reproduction Number

## Abstract

The recent West African Ebola virus pandemic, which affected >28,000 individuals increased interest in anti-Ebolavirus vaccination programs. Here, we systematically analyzed the requirements for a prophylactic vaccination program based on the basic reproductive number (R_0_, i.e. the number of secondary cases that result from an individual infection). Published R_0_ values were determined by a systematic literature research and ranged from 0.37 to 20. R_0_s ≥4 realistically reflected the critical early outbreak phases and superspreading events. Based on the R_0_, the herd immunity threshold (I_c_) was calculated using the equation Ic=1–(1/R_0_). The critical vaccination coverage (V_c_) needed to provide herd immunity was determined by including the vaccine effectiveness (E) using the equation Vc=Ic/E. At an R_0_ of 4, the I_c_ is 75% and at an E of 90%, more than 80% of a population need to be vaccinated to establish herd immunity. Such vaccination rates are currently unrealistic because of resistance against vaccinations, financial/ logistical challenges, and a lack of vaccines that provide long-term protection against all human-pathogenic Ebolaviruses. Hence, outbreak management will for the foreseeable future depend on surveillance and case isolation. Clinical vaccine candidates are only available for Ebola viruses. Their use will need to be focused on health care workers, potentially in combination with ring vaccination approaches.

## Introduction

Four Ebolaviruses (Ebola virus, Sudan virus, Bundybugyo virus, Taï Forrest virus) are endemic to Africa and can cause severe disease associated humans (1). Reston viruses are endemic to Asia and considered to be non-pathogenic in humans (1). However, very few genetic changes may result in human-pathogenic Reston viruses (1–3). Since the discovery of the first two members of the *Ebolavirus* family in 1976 in Sudan (today South Sudan) and Zaïre (today Democratic Republic of Congo), Ebolaviruses had until 2013 only caused small outbreaks in humans affecting up to a few hundred individuals (4,5). The recent Ebola virus outbreak in West Africa (2013–2016) resulted in 28,616 confirmed, probable, and suspected cases of Ebola virus disease and 11,310 deaths (5), which may still underestimate the actual numbers (6). It was the first Ebolavirus outbreak that affected multiple countries, was introduced to another country via air travel, and resulted in disease cases outside of Africa (4,5). The outbreak emphasized the health threats posed by Ebolaviruses and the importance of protection strategies (5,6).

Vaccination programs are effective in controlling infectious diseases, as demonstrated by the WHO-driven smallpox eradication (7). However, eradication seems to be an unrealistic aim for zoonotic viruses like the Ebolaviruses that circulate in animal reservoirs (8). Only herd immunity could prevent future outbreaks and protect individuals that cannot be vaccinated due to health issues (7). The herd immunity threshold (l_c_) describes the number of society members that need to be protected (l) to prevent outbreaks. It is based on the basic reproductive number R_0_ (number of secondary cases caused per primary case) of a pathogen (9–13).

Here, we performed a systematic analysis to determine the critical vaccine coverage (V_c_) required to prevent Ebolavirus outbreaks by a prophylactic mass vaccination program based on the R_0_ associated with Ebolavirus infection in humans. The results were further critically considered in the context of 1) the status of current Ebolavirus vaccine candidates and 2) the feasibility of a large-scale prophylactic Ebolavirus vaccination program taking into account a) the preparedness to participate in vaccination programs in the affected societies, b) logistic challenges, and c) costs.

## Methods

### Ebolavirus nomenclature

The nomenclature in this manuscript follows the recommendations of Kuhn et al. (14). The genus is *Ebolavirus*. The species are *Zaire ebolavirus* (type virus: Ebola virus), *Sudan ebolavirus* (type virus: Sudan virus), *Bundibugyo ebolavirus* (type virus: Bundibugyo virus), and *Taϊ Forest ebolavirus* (type virus: Taϊ Forest virus).

### Identification of studies that report on the basic reproductive number (R_0_) of Ebolaviruses

To identify scientific articles that have calculated the basic reproductive number (R_0_) for Ebolaviruses, we performed a literature search using PubMed (www.ncbi.nlm.nih.gov/pubmed) for the search term combinations “Ebola R0”, “Ebola basic reproductive number”, and “Ebola basic reproduction number” (retrieved on 29^th^ September 2017).

### Determination of herd immunity thresholds and their implications for Ebolavirus diseases prevention strategies

Based on the basic reproductive number R_0_, i.e. the number of secondary cases that result from an individual infection, the herd immunity threshold (I_c_) was calculated using equation 1

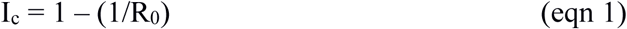

where I_c_ indicates the proportion of a society that needs to be protected from infection to achieve herd immunity. Next, the critical vaccination coverage (V_c_) that is needed to provide herd immunity was determined by including the vaccine effectiveness (E) using equation 2

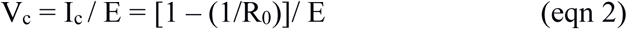

(9–13).

## Results

### Basic reproductive number (R_0_) values for Ebolaviruses

The PubMed search for “Ebola R0” provided 18 hits, the search for “Ebola basic reproductive number” provided 42 hits, and the search for “Ebola basic reproduction number” provided 35 hits (Figure 1; Data Sheet 1). After removal of the overlaps and inclusion of an additional article (identified from the reference list of (12)) this resulted in 51 articles, 35 of which provided relevant information on Ebolavirus R_0_ values (Figure 1; Data Sheet 1).

**Figure 1.**
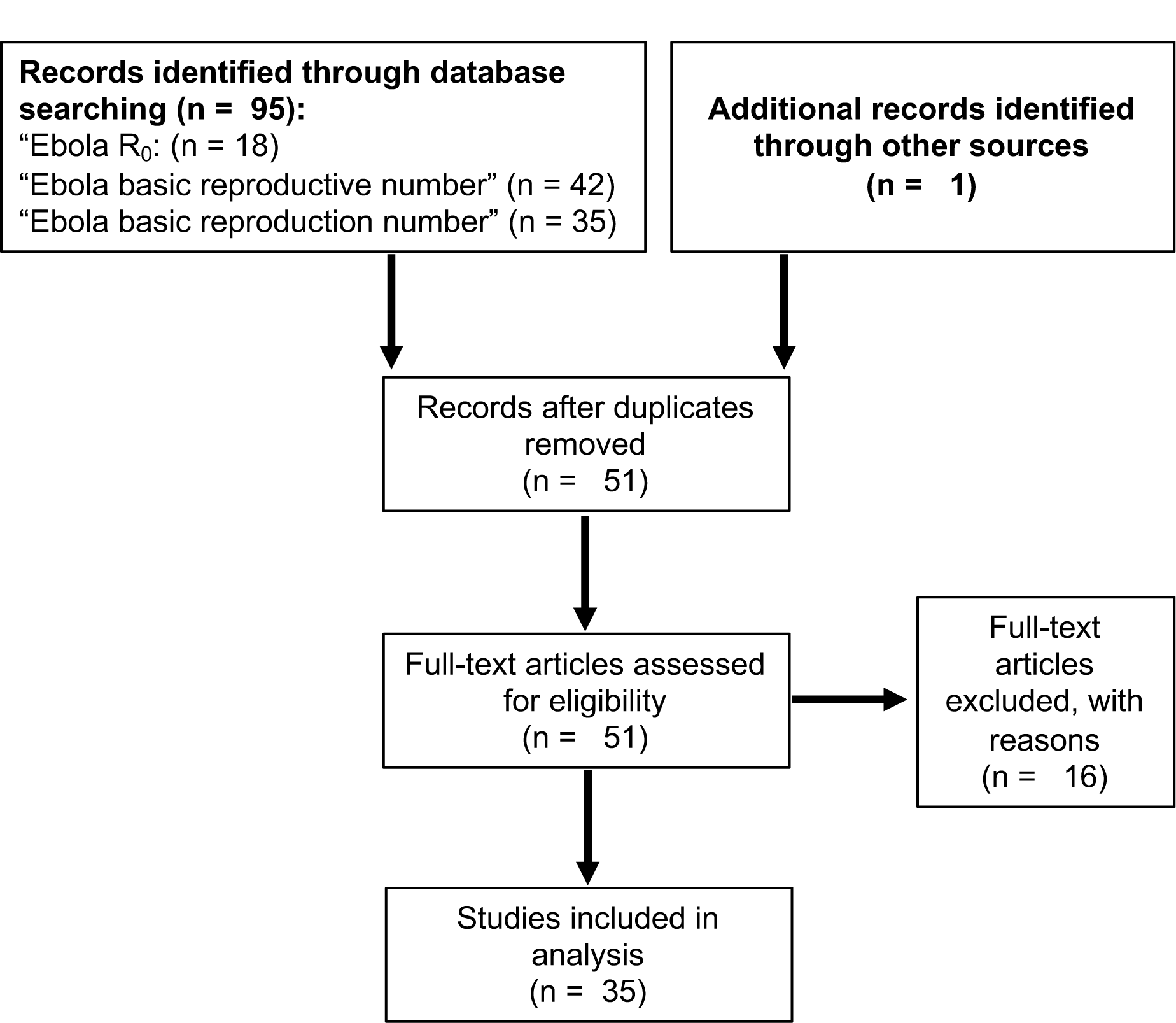
Summary of the literature search using PubMed (www.ncbi.nlm.nih.gov/pubmed) to identify articles that report on the basic reproductive number (R_0_) of Ebolaviruses.

R_0_ data were only available for Ebola virus and Sudan virus outbreaks. (Data Sheet 1). 29/35 studies analyzed data from the recent West African Ebola virus outbreak (Data Sheet 1). The others reported on Ebola virus outbreaks in the Democratic Republic of Congo. Four studies also included data from the Sudan virus outbreak 2000/2001 in Gulu, Uganda. We also considered a review that summarized all available data until February 2015 (4) (Data Sheet 1).

R_0_ indicates the number of new infections caused by an infected individual, and when greater than 1 an outbreak will spread. R_0_s ranged from 0.36 to 12 for Ebola virus and from 1.34 to 3.54 for Sudan virus (Data Sheet 1). Three studies directly compared the Ebola virus outbreak in Kikwit (1995, DR Congo) and the Sudan virus outbreak in Gulu (2000/ 2001, Uganda) (15–17), but did not reveal fundamental differences between the R_0_s (Data Sheet 1, Data Sheet 2).

Different approaches to calculate R_0_s lead to varying results (13). Accordantly, R_0_ values calculated for the Sudan virus outbreak 2000/ 2001 in Gulu using identical data ranged from 1.34 to 3.54 (Data Sheet 1, Data Sheet 2). Additionally, virus transmission is influenced by socio-economic and behavioral factors including the health care response, society perceptions, religious practices, population density, and/ or infrastructure (13,18). Concordantly, R_0_s determined in different districts of Guinea, Liberia, and Sierra Leone during the West African Ebola virus epidemic ranged from 0.36 to 3.37 (19). High reproductive numbers (≥4) are typically observed at the beginning of Ebolavirus outbreaks, prior to the implementation of control measures (20–23). Also, the spread of Ebolaviruses may be substantially driven by “superspreaders” who infect a high number (up to 15-20) of individuals (18,24-27). Hence, a vaccination program should establish herd immunity against Ebolaviruses that spread with an R_0_ of ≥4.

### Herd immunity threshold (Ic)

At an R_0_ of 4, the I_c_ (eqn 1) is 75%, which means that 75% of a population need to be immune to provide herd immunity (Figure 2A, Data Sheet 3). The I_c_ further rises to 80% at an R_0_ of 5, to 90% at an R_0_ of 10, and to 95% R_0_ of 20 (Figure 2A, Data Sheet 3). This shows that high proportions of a population need to be immune to establish effective herd immunity.

**Figure 2.**
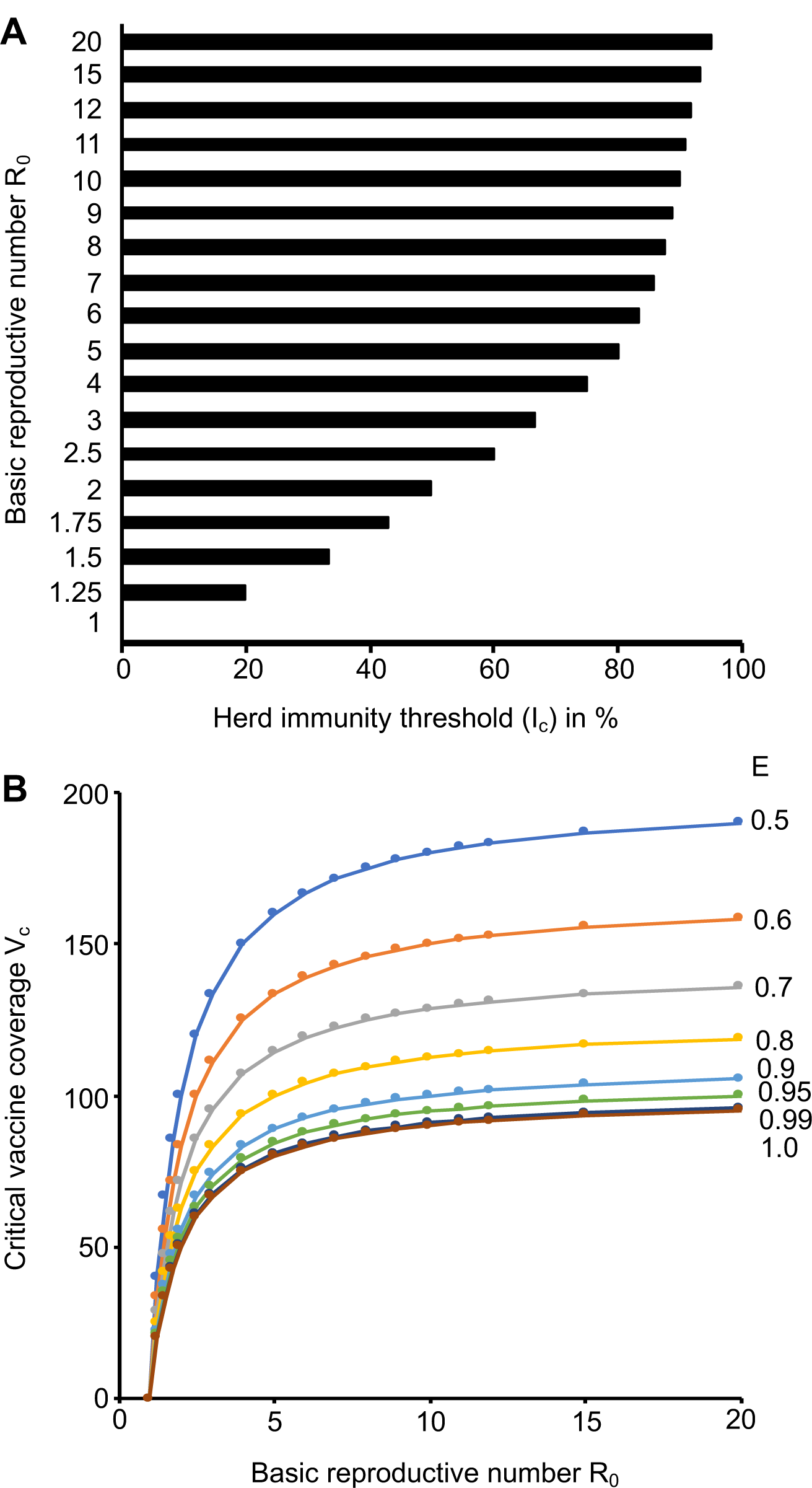
Herd immunity thresholds (Ic) and Critical vaccine coverage (Vc) values in dependence of the basic reproductive number (R_0_) and the vaccine efficacy (E). A) Ic values based on a range of R_0_ values that cover the range reported for Ebolaviruses. B) Vc values based on R_0_ values that cover the range reported for Ebolaviruses and E values that are in the range of those reported for approved vaccines. The respective numerical data are presented in Data Sheet 3.

### Critical vaccine coverage (V_c_)

As there is currently no approved vaccine for the prevention of Ebolavirus disease, we calculated a range of V_c_ (eqn 2) scenarios that reflect the efficacy range covered by approved vaccines. Attenuated replication-competent measles virus vaccines have been reported to protect up to 95% of individuals from disease after one dose, which increased to up to 99% after a second dose (28). The efficacy of varicella zoster virus vaccines, another attenuated replication-competent vaccine, was recently calculated to be 81.9% after one dose and 94.4% after two doses (29). Inactivated seasonal influenza virus split vaccines have been reported to have a substantially lower efficiency of 50-60% (30–32). Hence, we considered a V_c_ range between 50% and 100% (Figure 2B, Data Sheet 3). Vaccines, which provide high protection (ideally after a single vaccination), and high vaccination rates are required to prevent Ebolavirus outbreaks. If we assume an R_0_ of 4 and a vaccination efficacy E of 90%, more than 80% of a population need to be vaccinated to establish herd immunity. If the R_0_ rises to 5 a vaccine coverage of 80% would be required, even if a vaccine with 100% efficacy was available (Figure 2B, Data Sheet 3).

## Discussion

We performed an analysis of the Ebolavirus vaccine requirements to achieve the V_c_ needed for prophylactic mass vaccination programs. Vaccines need to prevent Ebolavirus outbreaks that spread at an R_0_≥4, which reflects the critical early outbreak phases prior to the implementation of control measures and superspreading events (18–27). At an R_0_ of 4, 80% of individuals need to be vaccinated with a vaccination efficacy of 90% to achieve herd immunity. Hence, highly effective vaccines and a high vaccination coverage are essential for successful prophylactic mass vaccination programs against Ebolaviruses.

Clinical vaccine candidates providing protection against all three to four human-pathogenic Ebolaviruses (Ebola virus, Sudan virus, Bundibugyo virus, potentially Taï Forest virus) do not currently exist (Data Sheet 4), although pre-clinical data suggest that the development of such vaccines may be feasible (5). Current vaccine candidates may also not provide the long-term protective immunity (≥10 years) necessary for sustainable protection against spillover events from animal reservoirs. Two studies reported immune responses 12 months after vaccination with different Ebola virus vaccine candidates (34,35). One of them described seroconversion in >90% of individuals after a single injection of rVSV-ZEBOV, a vesicular stomatitis virus-based Ebola virus vaccine. No or only a minor drop in antibody titers and neutralization capacity was reported 360 days after vaccination (34). A study investigating rVSV-ZEBOV and ChAd3-EBO-Z, a chimpanzee adenovirus type-3 vector-based Ebola virus vaccine, found lower seroconversion rates (rVSV-ZEBOV: 83.7%; ChAd3-EBO-Z: 70.8%) and reported the highest antibody response after one month and a decline afterwards (35). Thus, it is not clear, whether the vaccine-induced immunity covers the time frames >18 months that Ebolavirus survivors may remain contagious for (5,34,35,38–42). Notably, antibody responses may not always reliably indicate the protective efficacy of vaccines (36,37) (L. Lobel, unpublished data). Ebola virus recurrences and reinfections indicate that also natural Ebolavirus infections may not necessarily provide sustained protective immunity, which may further complicate the development of vaccines that provide long-term protection (43,44).

Limited acceptance of vaccinations may also limit Ebolavirus vaccination programs. In a rVSV-ZEBOV ring vaccination trial, only 5,837/ 11,841 patient contacts could be vaccinated. 34% of the contacts refused the vaccination (45). In a survey in Sierra Leone during the West African Ebola epidemic, 106/ 400 respondents (26.6%) were prepared to pay for a vaccination, while 290 respondents (72.5%) would have accepted a free vaccination (46). Hence, a I_c_ of 75% (necessary to prevent an outbreak that spreads with an R_0_=4 (Data Sheet 3)) would not be achievable, even under the threat of an ongoing epidemic.

The median maximum fee that survey participants in Sierra Leone during the West African Ebola epidemic were prepared to pay for a vaccine was about 5,000 leones ($0.65 as of 11th January 2018) (46). The international organization GAVI (www.gavi.org) is providing $5 million for the development of rVSV-ZEBOV, which is expected to pay for 300,000 vaccine doses (about $16.70/ dose) (47). Within a rVSV-ZEBOV ring vaccination trial, 11,841 contacts requiring vaccination from 117 clusters were identified over a ten-month period, i.e. about 101 individuals per confirmed Ebola virus disease patient (45). Hence, 300,000 doses will enable to vaccinate the contacts of approximately 2,970 Ebola virus disease patients. If an effective vaccine (which provided protection against all human-pathogenic Ebolaviruses) was available, a vaccination program would comprise about 462 million individuals in the countries that have been affected by Ebolavirus outbreaks (Data Sheet 5). Notably, the countries, which have been affected by Ebolavirus outbreaks so far, have large rural populations ranging from 13% (Gabon) to 84% (Uganda) (Data Sheet 5). Vaccination programs in rural areas are associated with logistical issues including transport difficulties, lack of equipment and trained medical specialists, and cultural and language barriers (48,49).

In conclusion, the achievement of a V_c_ of 75% that is necessary to prevent an outbreak that spreads with an R_0_ of 4 with a vaccine that has an efficacy of 100% is currently unrealistic because of limited vaccine acceptance in the affected populations and because of financial and logistical challenges. In addition, concurrent diseases such as HIV and cancer, along with potential side effects of vaccination, may remove significant numbers of potential vaccinees (5,50). Moreover, vaccines that provide long-term immunity against all three (or including Taï Forest virus, four) human-pathogenic Ebolaviruses, which would be needed to protect populations effectively from large Ebolavirus outbreaks in endemic areas, do not exist. Therefore, outbreak control of Ebolaviruses will for the foreseeable future depend on surveillance and the isolation of cases. Clinical vaccine candidates are only available for Ebola viruses and will need to be focused on health care workers, who are often involved in disease transmission (22), potentially in combination with the vaccination of patient contacts.

